# Genomic sequencing should extend to diverse priority pathogens for effective study and surveillance of antimicrobial resistance: a systematic review of whole-genome sequencing studies from India

**DOI:** 10.1101/2023.11.23.568416

**Authors:** Nazneen Gheewalla, Jaisri Jagannadham, Rintu Kutum, Shraddha Karve

**Affiliations:** Trivedi school of biology, Ashoka University, NH 44, Rajiv Gandhi Education City, Sonipat 131029, India; Centre for Bioinformatics and Computational Biology, Ashoka University, NH 44, Rajiv Gandhi Education City, Sonipat 131029, India; Department of computer science, Ashoka University, NH 44, Rajiv Gandhi Education City, Sonipat 131029, India

## Abstract

**Background:** Antimicrobial resistance (AMR) is a public health emergency in many low and middle-income countries, including India. To effectively tackle AMR, we need rapid diagnostics, effective surveillance and new antimicrobial drugs. Whole-genome sequencing of pathogens is the first definite step towards achieving these goals.

**Methods:** In this work, we review all the studies published till date that report whole-genome sequences of select priority AMR pathogens from India. We searched PubMed and Web of Science databases for the studies that involved whole-genome sequencing of AMR priority pathogens from India. For the top two highly sequenced pathogens, *S. typhi* and *K. pneumoniae*, we performed phylogenetic analyses to understand the geo-climatic distribution of genetically diverse strains.

**Results:** Our search reveals 94 studies that report 2547 unique whole-genome sequences. We find that most sequences are limited to select priority pathogens isolated from a couple of geo-climatic zones of India. Our phylogenetic analyses show that available data does not indicate systematic differences between the genomes of isolates from different geo-climatic zones. Our search also reveals complete absence of travel-related studies tracking possible movement of AMR pathogens within country. Lastly, we find very few studies that sequence AMR pathogens isolated from food, soil or other environments.

**Conclusion:** Together, these observations suggest that lndia should prioritize sequencing of diverse AMR pathogens from clinics as well as from environments and travellers rather than extending the geo-climatic range of already-sequenced pathogens. Our recommendations can be potentially valuable for other low and middle-income countries with limited resources, high prevalence of AMR and diverse geo-climatic conditions.

## Introduction

Antimicrobial resistance (AMR) is a global crisis and has been ranked amongst the top ten health concerns by WHO.^1^ In the year 2019, 1.27 million people died of infections by drug resistant pathogens.^2^ Antibiotic resistance is more prevalent in low and middle-income countries owing to lack of hygiene, resource limitation and other socio-economic factors.^3–5^ South Asia has the second largest burden of AMR, surpassed only by the sub-Saharan African region.^2^ Within South Asia, India is one of the hotspots for resistance. By the year 2050, global annual deaths due to infections by drug resistant pathogens are predicted to rise to ∼10 million with ∼2 million deaths in India alone.^4,6^ India also has a drug resistance index of 71, highest for any country in the world, indicating very poor efficacy for the existing drugs.^7^ AMR is thus a public health emergency for countries like India.

The diverse and dynamic nature of AMR demands continuous surveillance, rapid diagnostics and robust drug development programs, amongst other things. Traditionally, the first step of tackling resistant pathogen is antimicrobial susceptibility testing (AST). AST is a gold standard for determining resistance and is relatively economic to perform. However, AST has large turnaround times, requires customized workflow for every pathogenic group and cannot reveal the molecular mechanisms of resistance. As a result, the outcomes of ASTs are of little use for drug design or rapid diagnostics or extensive surveillance.

In the recent years, whole-genome sequencing (WGS) has been successfully used for understanding the molecular mechanisms of AMR,^8,9^ epidemiology of AMR^10,11^ as well as for drug discovery.^12^ WGS-based studies of antimicrobial resistant pathogens offer several advantages over other traditional approaches. First, WGS gives us a comprehensive picture of genomic changes in contrast to gene-based studies that are restricted to a few well-characterized genes.^13^ Second, WGS data can help establish the trajectories and timelines of resistance evolution.^14–17^ Third, WGS based approaches can potentially supplement or replace the traditional methods of AST.^13,18–22^ Fourth, current WGS technologies promise shorter turnaround times than traditional approaches.^23^ Short turnaround times are especially useful for slow-growing pathogens like *Mycobacterium*^24,25^ as well as in the treatment of conditions like septicemia where faster diagnosis can increase the patient survival rates significantly. Lastly, WGS-based approaches can also accurately identify pathogens that may have been misidentified by traditional methods. For instance, it is challenging to distinguish between *E. coli* and *Shigella* by methods other than whole-genome sequencing.^26^ These advantages in terms of diagnostics, therapeutics and surveillance automatically make genomic sequencing the first step in the fight against AMR and countries with high prevalence of AMR need a greater focus on genomic sequencing.^27^

To establish effective genomic sequencing for AMR in India, it is important to have information on the available whole-genome sequences from India. A comprehensive review of whole genome sequences of AMR pathogens from India is lacking. Sequence data available in most databases suggest that genomic sequencing from India is patchy at best. For example, according to the Bacterial and Viral Bioinformatics Resource Centre (BV-BRC),^28^ India is the third major contributor towards *S. typhi* sequences worldwide with the contribution of nearly 14%. But similar levels of sequencing efforts are lacking for almost all the other AMR pathogens. For instance, BV-BRC lists 8482 *E. coli* sequences from the USA as opposed to 827 sequences from India, which is four times as populated as the USA. Similarly, BV-BRC lists eight times as many of *K. pneumoniae* sequences from China than India.

Here we systematically review the studies that report whole-genome sequences of AMR pathogens from India. We focus on the pathogens that demand immediate attention according to the Indian Council of Medical Research.^29^ We find that the use of whole-genome sequencing in this context is very limited in India. Majority of the data are confined to a couple of geo-climatic zones within the country and is restricted to a few select pathogens. We then perform a phylogenetic analysis for the two pathogenic species that have been sequenced across most geo-climatic zones of India. Our results show that there are no systematic differences between the genomes of pathogens isolated from different geo-climatic zones (but see discussion for the limitations of the available data). This finding suggests that the immediate focus of sequencing efforts should be a number of diverse priority pathogens rather than already-sequenced pathogens from new geo-climatic zones of the country. We also find that the spread of resistance due to human travel remains severely understudied in the country. Lastly, our review shows that genomic studies of pathogens isolated from domestic animals, food and environment are rare in India. Increasing the sequencing efforts in this direction of ‘One health’^30^ can help gain a holistic understanding of resistance spread and evolution in India.

## Methods

### Choice of pathogenic species

We selected ten pathogenic species and one pathogenic genus for which immediate action is recommended.^29^ Specifically, this included eight species (*Escherichia coli, Enterobacter cloacae, Morganella morganii, Citrobacter koseri, Proteus mirabilis, Providencia rettgeri, Salmonella typhi, Serratia marcescens*) and one genus (*Klebsiella*) from the order *Enterobacterales*, and two other species from diverse taxa (*Acinetobacter baumannii* and *Candida auris)*. For the selected pathogens we included the genomic sequence of every isolate from AMR studies conducted in India, irrespective of the resistance profile of the pathogen. We reasoned that every sequence of a priority pathogen, resistant or sensitive to any specific antibiotic, is important for understanding the molecular mechanisms and evolution of resistance as well as for applications like surveillance and diagnostics.

### Literature Search

We searched the databases of PubMed and Web of Science on the 5 May, 2023 for studies that reported whole-genome sequences of selected AMR pathogens. We included only those studies that provided the data for pathogens isolated from Indian patients or from patients with a history of travel to India or environmental/food/veterinary samples collected in India. We followed PRISMA guidelines^31^ while performing the literature search. Our detailed search strategy is outlined in the supplementary material (Supplementary material 1).

### Checking for the uniform availability of whole-genome sequences for selected pathogens across different geo-climatic zones

To examine whether the number of available whole-genome sequences are uniform across different geo-climatic zones, we performed the Pearson’s chi square test of homogeneity using RStudio (v2022.07.0). To this end, we considered the pooled number of sequences of all the selected pathogens for each geo-climatic zone. We only used data for those isolates for which geo-climatic zone was known. To ascertain the geo-climatic zone of every isolate, we used the information reported in the corresponding study or the NCBI metadata associated with the genomic sequence. In case of a discrepancy in location mentioned in the study and NCBI metadata, we gave preference to the information provided in the relevant study.

### Phylogenetic Analysis of *S. typhi* and *K. pneumoniae*

To determine whether different lineages of a given organism are prevalent in different zones of India, we conducted a phylogenetic analysis. We chose *S. typhi* and *K. pneumoniae* for the phylogenetic analysis, as their genomic sequences were available from most geo-climatic zones of India. We excluded the sequences that lacked the information on the geo-climatic zone (Supplementary material 6), as well as the sequences where the accession numbers were not available.

We performed two separate phylogenetic analyses, one for 503 *S. typhi* genomic sequences and other for 231 *K. pneumoniae* genomic sequences. We first downloaded the genomic sequences from NCBI as complete genomes, raw reads or contigs. Using a custom script, we re-named all the downloaded sequences to denote their geo-climatic zone and the year of the study. We pre-processed all the raw reads, both single and paired-end, and checked the quality with FastQC v0.12.1.^32^ We then filtered the low-quality reads and adapters using Trimmomatic, v0.39.^33^ We used PhaME (Phylogenetics and Molecular evolution analysis tool, v1.0.2)^34^ for phylogenetic tree building. For the phylogenetic analysis of *S. typhi* sequences, we used *S. typhi CT18* as the outgroup reference.^35–38^ Similarly, we used *K. pneumoniae MGH78578* as an outgroup reference strain for the phylogenetic analysis of *K. pneumoniae sequences* and SPAdes genome assembler v3.15.5,^39^ for getting the draft genomes. We also performed a genotype analysis for *S. typhi* sequences using Genotyphi v2 scheme,^40^ and Kleborate v2.3.2^41^ for the genotype analysis of *K. pneumoniae* sequences. We used Iroki^42^ for visualization of the phylogenetic trees.

## Results

### Literature Search

The preliminary literature search for the selected pathogens yielded a total of 836 studies. We did not find any studies from India with WGS information on three species from our list, namely *Citrobacter koseri, Proteus mirabilis* and *Providencia rettgeri*. After exclusion of duplicates, we obtained 257 unique studies. From this subset, we further excluded 159 studies during the first screening based on the title, abstract and a brief view of full text wherever needed. Nine more studies were excluded during the second screening where we scrutinized the full text for the relevance to our review. This resulted in a total of 89 studies matching our inclusion criteria. We found 5 additional studies via back-referencing. In all we included 94 studies (that spanned over 9 years, from 2014 to 2023) in this systematic review that provided sequences and other data for 2547 isolates.

### Number of whole-genome sequences for any selected pathogen vary widely across different geo-climatic zones of India

The 94 studies that we shortlisted provide WGS data and related information for a total of 2547 distinct isolates. With 960 isolates, *S. typhi* represents a majority (37.69%) of these isolates (Figure 2). This is closely followed by the *Klebsiella* species which together form 31.99% of the isolates for which WGS data are available. For one of the Klebsiella species, *K. pneumoniae*, 811 sequences (31.84% of total sequences in our study) are available (Figure 2). We also obtained sequence data for 458 *E. coli* isolates, 222 *A. baumannii* isolates and 72 *C. auris* isolates. We pooled the sequence data for 17 *S. marcescens* isolates, 2 *E. cloacae* isolates and 1 isolate of *M. morganii*, 3 *K. quasipneumoniae* isolates and 1 *K. aerogenes* isolate during our analysis, (category ‘others’ in Figure 2) due to very low numbers of isolates from each group. Overall, these numbers show that the extent of whole-genome sequencing is highly variable for different pathogenic groups. *S. typhi* and *K. pneumoniae* are the most sequenced species among the selected pathogens, together comprising 69.53% of the total sequences. In contrast, we could not find even a single whole-genome sequence from India for three priority species from *Enterobacterales* – *P. rettgeri, P. mirabilis* and *C. koseri*.

**Figure 1.**
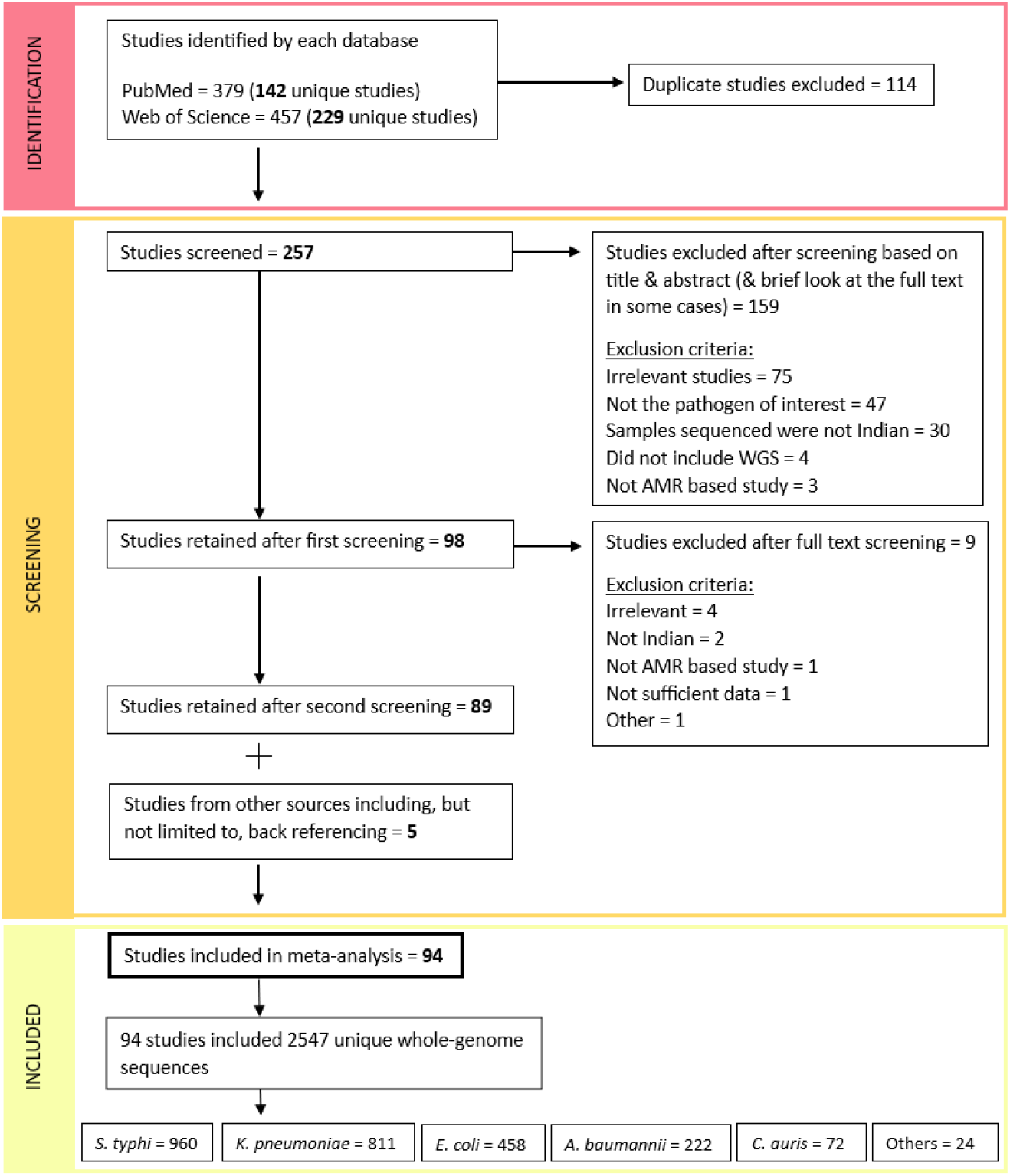
PRISMA flowchart describing the literature search strategy and outcomes for AMR related studies reporting whole–genome sequences of the selected pathogens. We searched PubMed and Web of Science databases for the relevant studies (see Supplementary material 1 for the search phrases). We screened 257 unique studies after removing duplicates within and between the search results from the two databases. We excluded 159 studies after screening the title and abstract. We further excluded 9 studies after the scrutiny of the full text. This resulted in 89 studies matching our inclusion criteria. We added 5 studies found by back-referencing or other sources. Together, we retained 94 studies for this systematic review. After removing the overlapping sequences across studies, the search yielded 2547 unique whole-genome sequences of the selected pathogens. The boxes in the bottom row show the number of genomic sequences for pathogenic species *S. typhi*, *K. pneumoniae*, *E. coli*, *A. baumannii* and *C. auris*. We found very few genomic sequences of *K. quasipneumoniae, S. marcescens, K. aerogenes, E. cloacae, M. morganii* that are clubbed together as ‘Others’.

**Figure 2:**
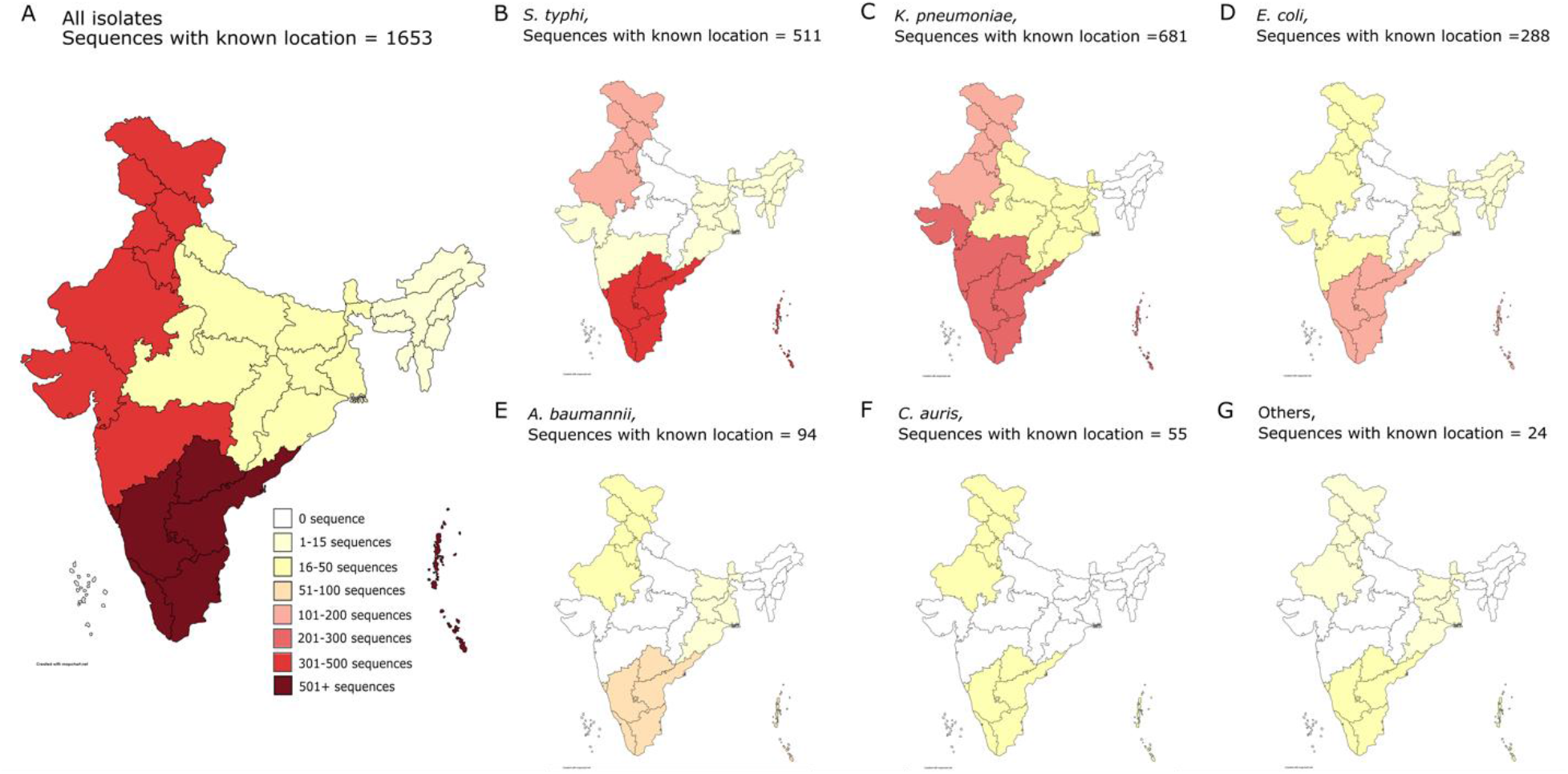
Number of whole-genome sequences of selected pathogens from every geo-climatic zone of India. **A.** The geo-climatic distribution of the all the 1653 whole-genome sequences with known locations. Panels **B-G** depict the distributions of number of isolates for individual pathogenic groups of *S. typhi, K. pneumoniae, E. coli, A. baumannii, C. auris* and others respectively. The group ‘others’ includes the species that have very few whole genome sequences, namely, *S. marcescens, K. quasipneumoniae, K. aerogenes, E. cloacae* and *M. morganii*. Only the isolates with known location are included. The colour of each geo-climatic zone in every panel (A-G) represents the range for the number of sequences as per the legend in the panel A. Exact number of sequences and their locations for each pathogenic group are in the supplementary materials 2 and 6 as well as in supplementary dataset. We used <u>mapchart.net</u> for the design and colour of the map, and <u>mapsofindia.com</u> as a reference for determining the geo-climatic zones of India.

We found only one study with *S. marcescens* where 17 isolates from patients and hospital environment had been sequenced.^130^ Similarly, for *E. cloacae* we found only two studies that reported the total of two whole genome sequences, while for *M. morganii and K. aerogenes*, we found one study with a single sequence each.^132–135^

We also looked at the geographic distribution of the whole-genome sequences across the six different geo-climatic zones of India (Figure 2). Out of the 2547 sequences, we could locate the geo-climatic zones of 1653 sequences (64.90% of the sequences) based on the information in the reported study or from the NCBI metadata associated with the genomic sequence. The number of sequenced genomes were highly variable across different geo-climatic zones for every selected pathogen (Figure 2). From the 1653 sequences with known locations, 893 (i.e. 54.02%) sequences were from the Southern zone, indicating that almost half the genomic sequences were from a single geo-climatic zone of India. Furthermore, out of these 893 genomic sequences from the southern zone, 581 sequences (i.e. more than 65%) belonged to *S*. *typhi* and *K. pneumoniae* (330 and 251 sequences respectively). These numbers indicate that the majority of the sequenced genomes from India were of two pathogenic species isolated from a single geo-climatic region.

Out of the remaining 760 genomic sequences from other geo-climatic zones, 362 were from the northern, 308 from western, 50 from eastern, 30 from central and only 10 were from the northeastern regions of India (Supplementary materials 2 and 6 and supplementary dataset). Formal statistical analysis validated our observation of large variation in the number of available whole-genome sequences across different geo-climatic zones for all the selected pathogens combined (Pearson’s chi square test for homogeneity, χ^2^= 502.25, *df* = 25, *p*<2.2×10^-16^). There is no evidence to indicate that the variation in extent of genomic sequencing reflects the population numbers or the disease burden in the respective geo-climatic zone.

The disparity in number of sequences is most likely the reflection of the disparity in the number of studies that report genomic sequences from each zone. There were only 2 studies which report genomic sequences from the northeastern zone as opposed to 43 studies that report WGS data from the southern zone. Similarly, we found only 2 studies from the central zone of India and both reported genomic sequences of *K. pneumoniae*. These numbers along with the other observations from our literature search suggest that often a particular research group focuses on a specific pathogen. A string of studies reporting WGS data of that pathogen then follows. Sequenced pathogenic isolates then typically belong to the same geo-climatic zone and often collected from the same locality.

In sum, genomic sequencing of pathogenic species is patchy in India and the current sequencing effort is focused on only a couple of pathogens mostly isolated from a couple of geo-climatic zones.

### Sequences of *S. typhi* from different geo-climatic zones of India do not form phylogenetically distinct clusters

Our literature review highlights that whole-genome sequencing of AMR pathogens in India needs to extend to all priority pathogens and cover more geo-climatic zones. But how different are the genomic sequences of isolates from different geo-climatic zones? The literature is equivocal on the sequence-level differences amongst isolates from different geo-climatic conditions with majority of the studies using geographic regions as a proxy for geo-climatic differences.^47,48,136^ Moreover, these studies tend to look at larger geo-climatic differences, often considering data across different countries and continents. Some studies suggest that different geographic regions may harbour genetically distant pathogens^47,48,136^ while others show genetic homogeneity in isolates from different geographic regions.^136^ Understandably, conclusions also vary for different pathogenic species.^136^ But little is known about the diversity of whole-genome sequences across different geo-climatic zones within a single country, like India.

To check whether the isolates from different geo-climatic zones of India form distinct sequence clusters we resorted to phylogenetic analysis of the available sequences. We chose the two priority pathogenic species, *S. typhi* and *K. pneumoniae*, which had isolates sequenced from most of the geo-climatic zones. Phylogenetic analysis of the 503 genomic sequences of *S. typhi* showed that the genomic sequences of isolates from the same geo-climatic zone do not cluster together (Figure 3). The conclusion is also supported by the phylogenetic analysis of 231 genomic sequences of *K. pneumoniae* (Supplementary material 3).

**Figure 3:**
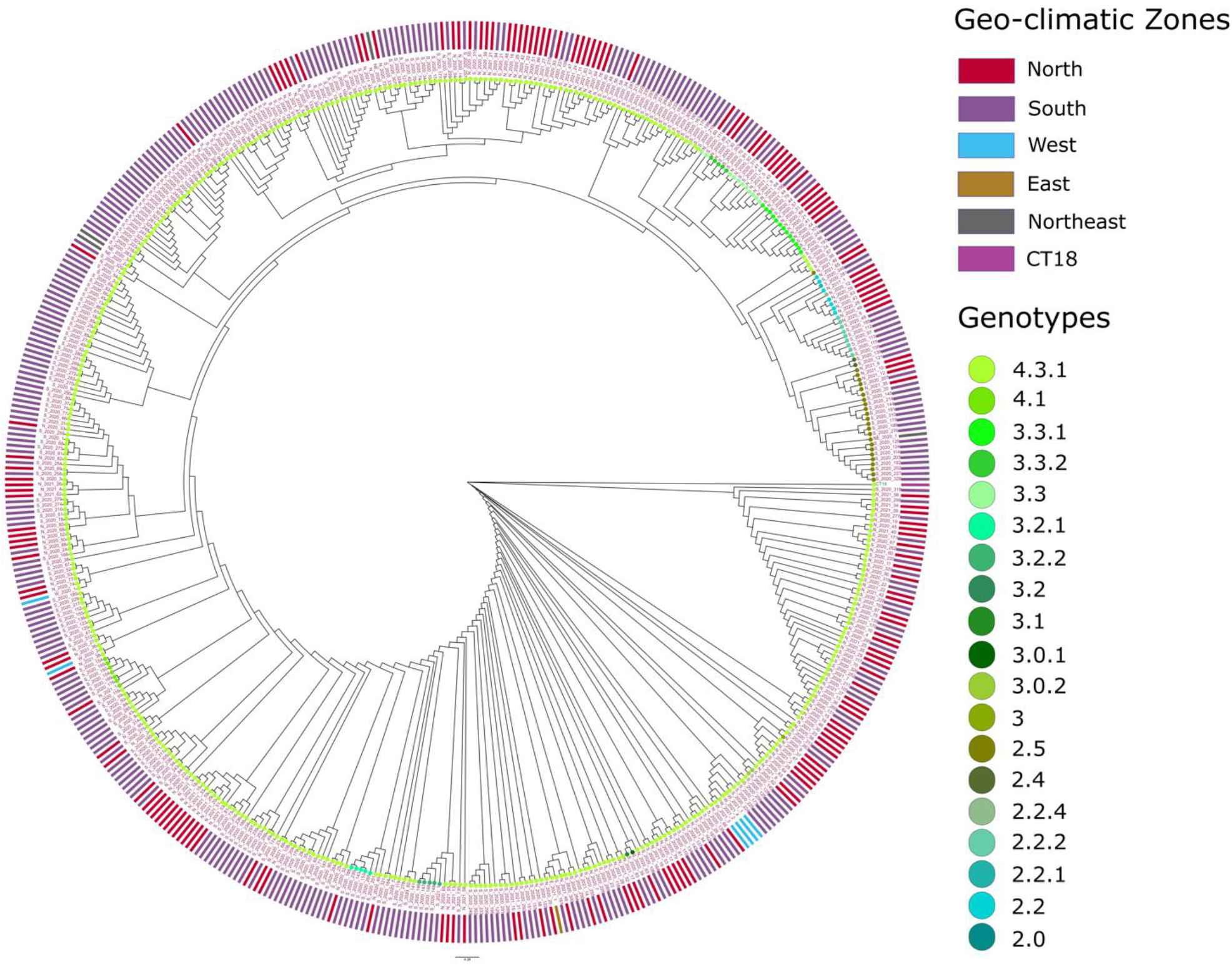
Phylogenetic analysis of *S. typhi* isolates. The geo-climatic zone wise distribution (outer circle) along with the genotype distribution (dots at end of leaf tip) and phylogenetic clustering of the 503 *S. typhi* isolates. The genotyping was done using Genotyphi v2, the phylogenetic tree was built using PhaMe v1.0.2, and visualization was created using Iroki. The name of the isolate indicates its geo-climatic zone and the year of study. Similar genotypes predominantly cluster together but not as per the geo-climatic zone of isolation (outer circle).

One possible reason for such homogeneity could be the spread of pathogens due to human dispersal across geo-climatic zones. The other reason could be the spread through environmental factors such as soil or water as well as food items. Therefore, we next checked for evidence for both these possibilities in the literature.

### Genomic sequencing of the pathogens from travel-associated infections is not common in India

Infected individuals traveling from one location to another can spread resistant strains of pathogens and tracking this movement is important to understand the spread of AMR.^137–139^ Most of the travel-related studies have a few limitations though. First, it is difficult to track the movement within the country, though some novel approaches may allow this.^140–142^ Second, majority of the travel-related studies do not take into account the acquisition of a pathogen prior to or during the travel.^94^ This may lead to incorrect identification of the source of infection as the travel destination. Third, most of these studies consider only symptomatic cases of the disease and asymptomatic carriers might be easily overlooked.^47,48,52,79,94,115,123^ Despite these limitations however, routine sequencing of isolates from the infected travellers can give a better picture of disease dispersal.^94,115^

Unfortunately, we found only one study that sequenced the genome of a pathogen possibly acquired during the travel between two geo-climatic zones of India.^44^ The patient, resident of the eastern zone, had a history of travel to the northern zone, but no clear link between the infection and travel could be established in the study. We found another handful of studies that sequenced isolates from the travellers that were visiting India/South Asia (Table 1 and Supplementary material 4). These studies identified the strain of the pathogen and established an association between the strain and specific geographic region. For example, Ingle et al. observed that *S. typhi* sub-lineage 4.3.1.2. was mainly associated with travellers returning from India and carried mutations in the quinolone resistance-determining genomic regions.^47,48^ Similarly, Yaita et al. studied the acquisition of ESBL-producing *E. coli* by travellers returning to Japan. The study found that the majority of the people returning from India (10 out of 14) tested positive for ESBL-producing *E. coli.*^94^ Yet another study found that an individual returning to Switzerland from India was colonised with a carbapenamase-producing *E. coli* harbouring a blaOXA-484, a variant of carbapenemase new to Switzerland.^115^

**Table 1:**
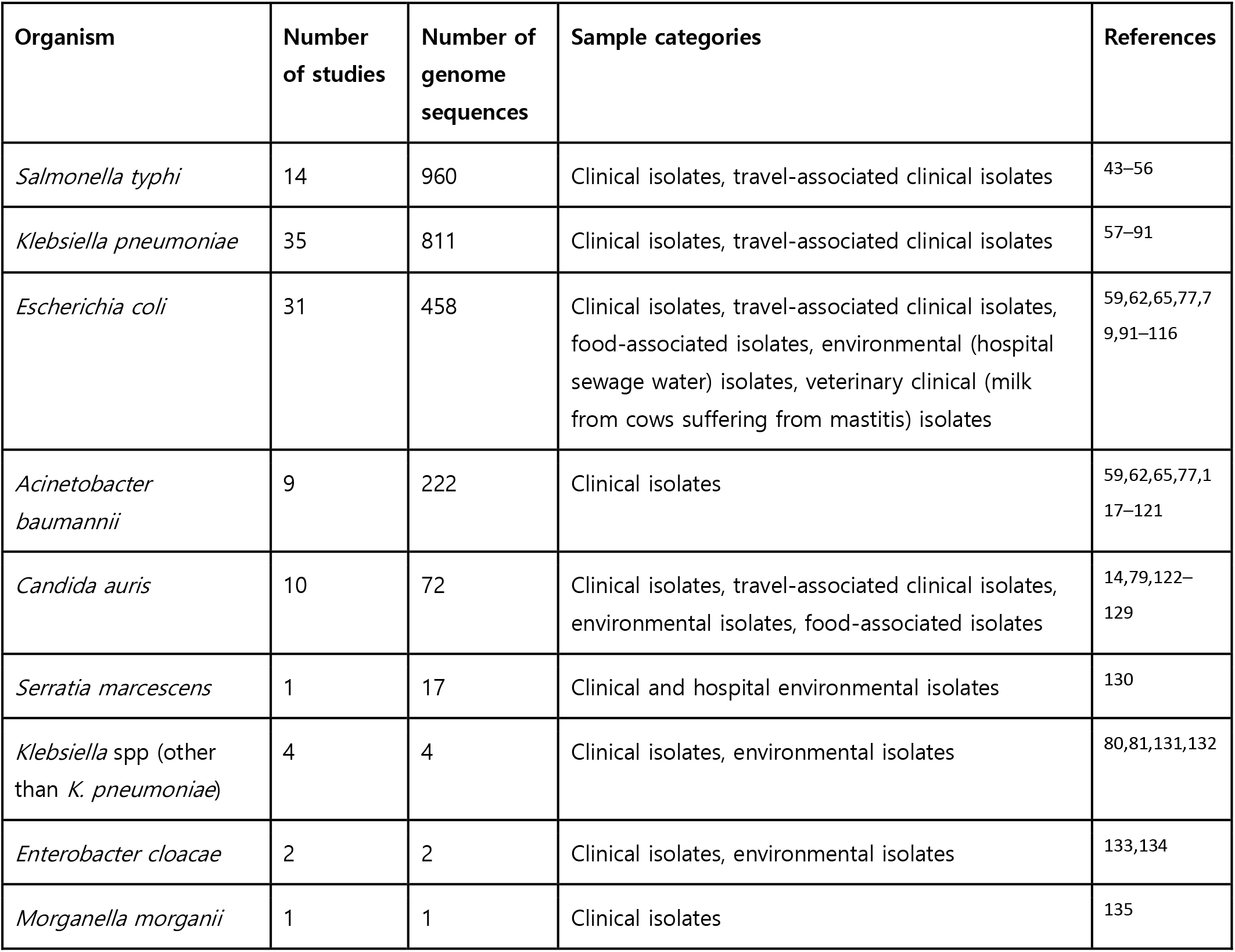
Number of studies and number of unique whole-genome sequences therein for the selected pathogenic groups. The column ‘Sample category’ indicates whether the isolates were from patients from India (clinical) or from patients with a travel history to India (travel-associated clinical) or from environmental/ food/ veterinary samples collected in India.

| Organism | Number of studies | Number of genome sequences | Sample categories | References |
| --- | --- | --- | --- | --- |
| *Salmonella typhi* | 14 | 960 | Clinical isolates, travel-associated clinical isolates | 43–56 |
| *Klebsiella pneumoniae* | 35 | 811 | Clinical isolates, travel-associated clinical isolates | 57–91 |
| *Escherichia coli* | 31 | 458 | Clinical isolates, travel-associated clinical isolates, food-associated isolates, environmental (hospital sewage water) isolates, veterinary clinical (milk from cows suffering from mastitis) isolates | 59,62,65,77,79,91–116 |
| *Acinetobacter baumannii* | 9 | 222 | Clinical isolates | 59,62,65,77,117–121 |
| *Candida auris* | 10 | 72 | Clinical isolates, travel-associated clinical isolates, environmental isolates, food-associated isolates | 14,79,122–129 |
| *Serratia marcescens* | 1 | 17 | Clinical and hospital environmental isolates | 130 |
| *Klebsiella* spp (other than *K. pneumoniae*) | 4 | 4 | Clinical isolates, environmental isolates | 80,81,131,132 |
| *Enterobacter cloacae* | 2 | 2 | Clinical isolates, environmental isolates | 133,134 |
| *Morganella morganii* | 1 | 1 | Clinical isolates | 135 |

These examples illustrate the effective tracking of the resistance spread from India to other countries. We did not find similar studies for individuals returning to India from other regions of the world or for individuals traveling across different geo-climatic zones of India. These observations demand increased genomic sequencing of pathogens isolated from travellers to better understand the resistance dynamics within the country.

### Whole-genome sequencing of AMR isolates from food and environment is limited in India

We next looked for the genomic sequences of priority pathogens isolated from environmental and food samples. AMR studies from all over the world have reported widespread presence of resistance genes in water, soil as well as in food items.^143–145^ This ‘One health’ approach of study underlines the importance of tracking resistance across abiotic environments alongside the clinics.^30^ We thus looked for studies that reported whole-genome sequences of priority pathogens isolated from environments or food items.

Our search uncovered 3 such studies of environmental samples from India. It was striking that even such handful of studies (Supplementary material 5) reported isolates that harbour antibiotic resistant markers.^124,132,134^ For example, an isolate of *K. aerogenes* from agricultural soil was found to contain ∼30 AMR genes.^132^ Another sequencing study demonstrated that 23 (out of 24) *C. auris* isolates from marshy wetlands and sandy beach areas of Andaman and Nicobar Islands were resistant to a variety of antifungals.^124^

We also found 5 studies from India that reported sequences of AMR pathogens isolated from food sources (Supplementary material 5).^93,101,104,109,122^ For instance, a study reported resistance patterns in the yeast species found on the surface of apples.^122^ Authors found that 73% of the total isolates belonged to *Candida spp.* and 11% of those isolates were of the priority species *C. auris*. Alarmingly, most of these isolates (15 out of 16) had heightened resistance against fluconazole.^122^ Similarly, multiple sequencing studies found ESBL producing *E. coli* from poultry samples and one group demonstrated that 78.5% of their poultry isolates were multi-drug resistant.^101,104^ Another sequencing study from the Banaskantha district of Gujarat reported that 90% of their samples procured from ∼30 different farms contained *E. coli*. Nearly 80% of these *E. coli* were ESBL producers.^109^ There were also a few studies reporting drug-resistant *E. coli* from bovine mastitis raising a concern of pathogens getting into the milk and milk products.^111,113,114^

Overall the results indicate a high prevalence of AMR priority pathogens in the environments and food items but extensive and systematic sequencing studies are needed to obtain a truly comprehensive picture.

## Discussion

Our literature search revealed that whole-genome sequencing of AMR priority pathogens from India is patchy, both in terms of species that were sequenced and geo-climatic zones where the pathogens were isolated. For three of the priority pathogens *Citrobacter koseri, Proteus mirabilis, Providencia rettgeri* we did not find even a single genomic sequence reported from India. Recent evidence suggests that all the three species are clinically important.^146–152^ For example, multi-drug and pan-drug resistant varieties of *P. rettgeri* caused nosocomial infections in India.^148^ Similarly, a longitudinal study from a tertiary healthcare centre in western India reported *C. koseri* with beta-lactam and carbapenem resistance over a period of 3 years.^149^ Another study found a nosocomial NICU outbreak of multi-drug resistant *P. mirabilis* that resulted in 80% mortality.^150^ Yet another study found ESBL positive *P. mirabilis* in food and livestock samples.^153^ Apart from these three pathogens that lack any WGS data from India, we also found other important pathogenic groups that were severely underrepresented. For instance, carbapenem-resistant *Acinetobacter* is a serious threat and leading cause of nosocomial infections in India^29^ and yet we found only 222 whole-genome sequences of *Acinetobacter baumannii* from the whole country.

We performed phylogenetic analyses on the sequence data of two highly-sequenced priority pathogens *S. typhi* and *K. pneumoniae*. Results showed that genomic sequences of isolates from different geo-climatic zones do not form distinct clusters (Fig 3, Supplementary material 3). In the absence of evidence for distinct sequence diversity across different geo-climatic zones, it might be beneficial to prioritize the sequencing of diverse pathogenic species over the increased coverage of already-sequenced species across different geo-climatic zones of India. However, there are some limitations of the datasets that we used for our phylogenetic analysis, as discussed below.

Our conclusions are drawn from the limited sequence data that are available from India. The sequences used in the phylogenetic analysis were collected over a period of nine years. Insufficient data for individual years prevented us from performing the analysis separately for each year. Additionally, in many cases there was no clear indication of the month and year of isolation. Adding this information to the phylogenetic analysis may change the inferences drawn. It is reasonable to assume that a particular resistant variant of a pathogen is not likely to arise (or arrive, with an infected individual/food item/environmental agent) in multiple geo-climatic zones at the same time. But how rapidly any resistant variant spreads can be dependent on several factors such as mode of transmission, fitness advantage over the prevalent variants and climatic conditions. Certain resistant variants of any pathogen might already be widespread across all geo-climatic zones. Systematic and continued genome sequencing across all geo-climatic zones is needed to resolve these possibilities for every priority pathogen. It will also help to have a clearer indication of time of isolation and geo-climatic zone for all the reported sequences. For example, from the existing literature, we could not find the geo-climatic zone for 449 sequences of *S. typhi* and adding this information may change the inferences of the phylogenetic analysis.

We also note that for the phylogenetic analyses we used two gram-negative enteric pathogens, *S. typhi* (Figure 3) and *K. pneumoniae* (Supplementary material 3). Genomic sequences of other non-enteric or gram-positive pathogenic species may cluster according to the geo-climatic zones. It is also known that the ratio of core to accessory (or dispensable) genes varies across different pathogenic species.^154^ Such variation can affect the conclusions drawn from phylogenetic and MLST analysis.^155^ The predominant nature of the infections, nosocomial vs community-based, may also affect the spread across geo-climatic zones. For instance, *Acinetobacter baumannii* is primarily a nosocomial pathogen^156,157^ and specific strains may dominate specifies geo-climatic regions.

At the time of our literature search, only two geo-climatic zones had reasonable representation for *S. typhi* (330 and 170 sequences each from Southern and Northern zones) while remaining zones of West, Northeast, East had reported 6, 4 and 1 genomic sequence respectively. But genomic sequences from even Southern and Northern zones did not cluster separately. A possible reason for this could be that most of the reported genomic sequences (408 out of 503) belonged to the *S. typhi* lineage 4.3.1, suggesting one dominant infectious strain throughout the country. This may not be the case with other important pathogens and may lead to different observations. However, another study^85^, as well as our own analysis with *K. pneumoniae* (Supplementary material 3) genomes, demonstrates lack of distinct clusters for genomic sequences from different geo-climatic regions.

We reasoned that homogeneity in genomic sequences across different geo-climatic zones might be due to the spread of infectious pathogens/resistance genes through human travellers or food items or other environments. Documentation and sequencing of such instances is important as it can allow effective mapping of the isolates’ ancestry and help understand their dispersal patterns. Sequencing the isolates from infected travellers can also uncover the original incidences of certain infections. For example, a recent study discovered that the first incidence of *C. auris* infection was a 54-year-old male travelling to India in 2007, which was two years before the supposed first case of *C. auris* infection was reported from Japan.^123^ This discovery has altered the timeline of emergence of *C. auris* infections.

Our literature search revealed very few studies with genomic sequences of pathogens from travel-related infections. Majority of the studies were travellers who acquired the infection in India and travelled abroad. There were almost no studies reporting sequences of pathogens acquired during the incoming travel to India or more importantly, travel within the country. This observation underlines the necessity of tracking travellers for possible infections and extending these investigations to include whole-genome sequencing of the pathogen.

Whole-genome sequences of isolates from soil, water, other environments and foods were also modest in number. Alarmingly, even these few reports uncovered important resistance markers. For example, we found studies that reported colistin-resistant pathogens from food samples. Colistin is one of the final-resort antibiotics and the presence of colistin resistant *Enterobacteriaceae* (group3 pathogens as per ICMR)^29^ in food items is concerning. Colistin-resistant *Enterobacteriaceae* (like *Klebsiella, Enterobacter, Citrobacter, E. coli*) and *Pseudomonas* were found in a range of food samples collected from shops and households ^93^. Moreover, few of these isolates contained the *mcr-1* gene which is responsible for plasmid-mediated spread of colistin resistance. Extrinsic resistance elements, such as plasmids, can spread the resistance rapidly across pathogens from different groups.^158^ Our results underline the need for comprehensive ‘One health’ approach in WGS studies with extensive sequencing of isolates from environments.

## Conclusion

Our review collates the studies that sequence the genomes of priority AMR pathogens from India. We find that many priority pathogens are not routinely sequenced in India while some have not been sequenced at all. Additionally, most genomic sequences are available from only a couple of geo-climatic zones. With the limited sequence data that is available, we infer that genomic sequence diversity is homogeneous across the geo-climatic zones. To assert or refute this conclusion however, we need systematic genomic sequencing for a few more priority pathogens across all geo-climatic zones. This task is resource-intensive and implementation may take a few years at least. While such comprehensive sequencing data is being generated across the country we need to urgently begin sequencing diverse priority pathogens such as *A. baumannii*, *C. koseri, P. mirabilis* and *P. rettgeri*. This should be accompanied by genomic sequencing of isolates from travel-related infections and environment. Our recommendations can be valuable for other low and middle-income countries with diverse geo-climatic conditions, high prevalence of AMR and limited resources.

## Acknowledgements

This project has received funding from the Rockefeller Foundation, Mphasis M1 Foundation and Axis Bank. We thank Dr. Srikrishna Subramanian for insightful discussions and Dr. Magdalena San Roman as well as Dr. Abhishek Mishra for comments on the manuscript draft. We also thank Dr. Punit Kaur and Dr. Mohit Bhatia for providing sample accession numbers and metadata on request. We thank Centre for Bioinformatics & Computational Biology supported by Department of Biotechnology, for providing computing resources for running the programs/scripts for phylogenetic analysis.

## Author contributions

S.K and N.G. conceptualised the study. N.G. carried out the literature review and data screening. N.G and S.K were involved in the analysis, wrote and edited the paper. J.J performed the phylogenetic analysis and commented on the draft of the manuscript. R.K helped with the statistical analysis and phylogenetic analysis and commented on the draft of the manuscript.

