## supplementary materials 1-6 for "Genomic sequencing should extend to diverse priority pathogens for effective study and surveillance of antimicrobial resistance: a systematic review of whole-genome sequencing studies from India"

**Supplementary material**

**Supplementary material 1:**

**Details of the literature review:**

We conducted extensive search for antimicrobial resistance (AMR) related studies including whole-genome sequences (WGS) for the select pathogens of interest using PubMed and Web of Science databases. We used following search phrases for this purpose.

For PubMed

1. ((Pathogen of interest*) AND (Whole Genome Sequencing)) AND (India)
2. (((((Antibiotic Resistance) OR (AMR)) OR (Antimicrobial Resistance)) AND (Pathogen of interest*)) AND (Whole Genome Sequencing)) AND (India)

For Web of Science

1. Pathogen of interest* AND Whole Genome Sequencing AND India
2. (Antibiotic Resistance OR AMR OR Antimicrobial Resistance) AND (Pathogen of interest* AND Whole Genome Sequencing AND India)

* We used the name of each pathogen/pathogenic group separately.

**Supplementary material 2:**

**Description of supplementary dataset:**

The dataset lists 94 studies included in this systematic review. The dataset is organized as per pathogen or pathogenic group. So a study that sequenced multiple pathogens or pathogenic groups of interest will be repeated in this dataset. For each of the chosen pathogen we list the studies which provide WGS data and related metadata. The dataset gives the study title, number of unique sequences found in the study as well as the zone and source of the sample (clinical/clinical-environmental/clinical-veterinary/environmental/food) wherever available.

**Supplementary figure 3:**

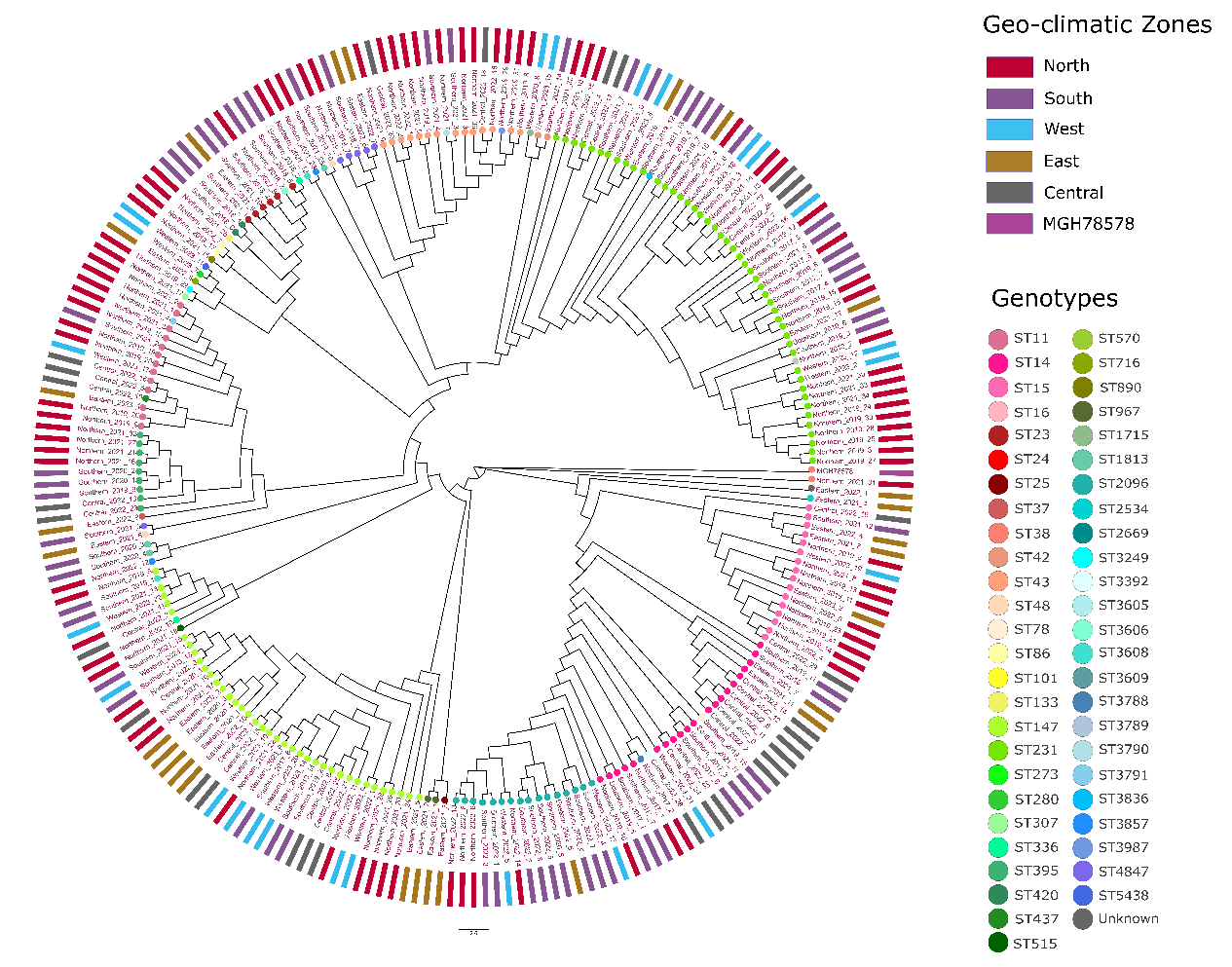

**Phylogenetic analysis of 231 *K. pneumoniae* isolates.** The geo-climatic zone wise distribution (outer circle) along with the MLST (dots at end of leaf tips) and phylogenetic clustering of the 231 *K. pneumoniae* isolates. The MLST analysis was done using Kleborate v2.3.2, the phylogenetic tree was built using PhaMe v1.0.2 and *K. pneumoniae* MGH78578 was used as an outgroup. The tree was visualised using Iroki. The name of the isolate indicates its zone and the year of study. Similar genotypes predominantly cluster together but not as per the geo-climatic zone of isolation (outer circle).

**Supplementary material 4:**

**Details of travel associated studies found in our literature search:**

Supplementary material 4 describes the travel related studies from the 94 studies shortlisted for the metaanalysis.

| **Sr No** | **Organism** | **Title** | **Number of unique isolates with WGS** | **Location** | **Zone** | **Sample source** | **Reference** |
| --- | --- | --- | --- | --- | --- | --- | --- |
| 1 | *C. auris* | Earliest case of Candida auris infection imported in 2007 in Europe from India prior to the 2009 description in Japan | 1 | Unknown | Unknown | Clinical | ^1^ |
| 2 | *C. auris* | Simultaneous Infection with Enterobacteriaceae and Pseudomonas aeruginosa Harboring Multiple Carbapenemases in a Returning Traveler Colonized with Candida auris | 1 | Unknown | Unknown | Clinical | ^2^ |
| 3 | *S. typhi* | Emergence of ceftriaxone resistant Salmonella enterica serovar Typhi in Eastern India | 1 | Kolkatta | East^#^ | Clinical | ^3^ |
| 4 | *S. typhi* | Informal genomic surveillance of regional distribution of Salmonella Typhi genotypes and antimicrobial resistance via returning travellers | 191 | Unknown | Unknown | Clinical | ^4^ |
| 5 | *S. typhi* | Genomic Epidemiology and Antimicrobial Resistance Mechanisms of Imported Typhoid in Australia | 87 | Unknown | Unknown | Clinical | ^5^ |
| 6 | *S. typhi* | Emerging high-level ciprofloxacin-resistant Salmonella enterica serovar typhi haplotype H58 in travelers returning to the Republic of Korea from India | 8 | Korean patients with history of travel to Northwest India | North* | Clinical | ^6^ |
| 7 | *K. pneumoniae* | Simultaneous Infection with Enterobacteriaceae and Pseudomonas aeruginosa Harboring Multiple Carbapenemases in a Returning Traveler Colonized with Candida auris | 1 | Unknown | Unknown | Clinical | ^2^ |
| 8 | *E. coli* | Epidemiology of extended-spectrum β-lactamase producing Escherichia coli in the stools of returning Japanese travelers, and the risk factors for colonization | 14 | Unknown | Unknown | Clinical | ^7^ |
| 9 | *E. coli* | Simultaneous Infection with Enterobacteriaceae and Pseudomonas aeruginosa Harboring Multiple Carbapenemases in a Returning Traveler Colonized with Candida auris | 1 | Unknown | Unknown | Clinical | ^2^ |
| 10 | *E. coli* | Repatriation of a patient with COVID-19 contributed to the importation of an emerging carbapenemase producer | 1 | Unknown | Unknown | Clinical | ^8^ |

\# The study reports a travel history to North India but does not make any further connection on acquisition of infection during travel. Hence we consider the sample to be from the Eastern zone. This is the only travel associated sample included in the phylogenetic analysis of *S. typhi* as the travel was within the country.

* Despite of a travel history to Northwest as per the study we placed the isolates under the Northern Zone as most of the travel was in the North zone.

**Supplementary table 5:**

**Studies that report the WGS data from food and environmental samples from India.** The table lists the studies which reported whole-genome sequences of isolates from food or environmental samples.

| **Sr No** | **Organism** | **Title** | **Number of unique isolates with WGS** | **Location** | **Zone** | **Sample source** | **Reference** |
| --- | --- | --- | --- | --- | --- | --- | --- |
| 1 | *C. auris* | Candida auris on Apples: Diversity and Clinical Significance | 16 | Delhi | North | Food | ^9^ |
| 2 | *C. auris* | Environmental Isolation of Candida auris from the Coastal Wetlands of Andaman Islands, India | 24 | Andaman & Nicobar Islands | South | Environmental | ^10^ |
| 3 | *E. cloacae* | Draft genome sequences of nonclinical and clinical Enterobacter cloacae isolates exhibiting multiple antibiotic resistance and virulence factors | 2 | Bhubaneshwar | East | 1 Environmental & 1 Clinical | ^11^ |
| 4 | *K. aerogenes* | Whole genome sequencing data of Klebsiella aerogenes isolated from agricultural soil of Haryana, India | 1 | Haryana | North | Environmental | ^12^ |
| 5 | *E. coli* | Detection of chromosomal and plasmid-mediated mechanisms of colistin resistance in Escherichia coli and Klebsiella pneumoniae from Indian food samples | 1 | Chennai | South | Food | ^13^ |
| 6 | *E. coli* | Risk of Transmission of Antimicrobial Resistant Escherichia coli from Commercial Broiler and Free-Range Retail Chicken in India | 10 | Hyderabad, Bellary, Mysore | South | Food – poultry | ^14^ |
| 7 | *E. coli* | Genomic and Functional Characterization of Poultry Escherichia coli From India Revealed Diverse Extended-Spectrum beta-Lactamase-Producing Lineages With Shared Virulence Profiles | 10 | Hyderabad, Bellary, Mysore | South | Food – poultry | ^15^ |
| 8 | *E. coli* | Whole genome sequencing and characteristics of extended-spectrum beta-lactamase producing Escherichia coli isolated from poultry farms in Banaskantha, India | 1 | Banaskantha | West | Food – poultry | ^16^ |

**Supplementary table 6:**

**Zone wise distribution of whole-genome sequences for each selected pathogen:** The table lists the number of whole-genome sequences from every geo-climatic zone of India for every selected pathogen. The penultimate column shows the number whole-genome sequences for that organism with unknown location.

| **Organism** | **South** | **North** | **East** | **West** | **Central** | **Northeast** | **Unknown** | **Total no of sequences** |
| --- | --- | --- | --- | --- | --- | --- | --- | --- |
| Others* | 21 | 1 | 2 | 0 | 0 | 0 | 0 | **24** |
| *C. auris* | 26 | 29 | 0 | 0 | 0 | 0 | 17 | **72** |
| *A. baumannii* | 68 | 21 | 5 | 0 | 0 | 0 | 128 | **222** |
| *E. coli* | 197 | 33 | 13 | 39 | 0 | 6 | 170 | **458** |
| *K. pneumoniae* | 251 | 108 | 29 | 263 | 30 | 0 | 130 | **811** |
| *S. typhi* | 330 | 170 | 1 | 6 | 0 | 4 | 449 | **960** |
| **TOTAL** | **893** | **362** | **50** | **308** | **30** | **10** | **894** | **2547** |

* The group ‘Others’ includes the pathogens that have very few whole genome sequences, namely, *S. marcescens, K. quasipneumoniae, K. aerogenes, E. cloacae* and *M. morganii*.
